# Trade-offs constrain adaptive pathways to T6 survival

**DOI:** 10.1101/2022.09.02.506412

**Authors:** Kathryn A. MacGillivray, Siu Lung Ng, Sophia Wiesenfeld, Randi L. Guest, Tahrima Jubery, Thomas J. Silhavy, William C. Ratcliff, Brian K. Hammer

## Abstract

Many microbial communities are characterized by intense competition for nutrients and space. One way for an organism to gain control of these resources is by eliminating nearby competitors. The Type VI Secretion System (T6) is a nano-harpoon used by many bacteria to inject toxins into neighboring cells. While much is understood about mechanisms of T6-mediated toxicity, little is known about the ways that competitors can defend themselves against this attack, especially in the absence of their own T6. Here we use directed evolution to examine the evolution of T6 resistance, subjecting eight replicate populations of *Escherichia coli* to T6 attack by *Vibrio cholerae*. Over ~500 generations of competition, the *E. coli* evolved to survive T6 attack an average of 27-fold better than their ancestor. Whole genome sequencing reveals extensive parallel evolution. In fact, we found only two pathways to increased T6 survival: *apaH* was mutated in six of the eight replicate populations, while the other two populations each had mutations in both *yejM* and *yjeP*. Synthetic reconstruction of individual and combined mutations demonstrate that *yejM* and *yjeP* are synergistic, with *yejM* requiring the mutation in *yejP* to provide a benefit. However, the mutations we identified are pleiotropic, reducing cellular growth rates, and increasing susceptibility to antibiotics and elevated pH. These trade-offs underlie the effectiveness of T6 as a bacterial weapon, and help us understand how the T6 shapes the evolution of bacterial interactions.

**Significance:** Bacteria are the most abundant organisms on Earth and often live in dense, diverse communities, where they interact with each other. One of the most common interactions is antagonism. While most research has focused on diffusible toxins (e.g., antibiotics), bacteria have also evolved a contact-dependent nano-harpoon, the Type VI Secretion System (T6), to kill neighboring cells and compete for resources. While the co-evolutionary dynamics of antibiotic exposure is well understood, no prior work has examined how targets of T6 evolve resistance. Here, we use experimental evolution to observe how an *Escherichia coli* target evolves resistance to T6 when it is repeatedly competing with a *Vibrio cholerae* killer. After 30 rounds of competition, we identified mutations in three genes that improve *E. coli* survival, but found that these mutations come at a cost to other key fitness components. Our findings provide new insight into how contact-dependent antagonistic interaction drives evolution in a polymicrobial community.

## Introduction

Bacteria are one of the most common forms of life on Earth and often live in polymicrobial biofilms. Within this complex community, negative bacterial interactions are the norm^1^, constantly competing for resources such as nutrients and space. One way for bacteria to gain an advantage over their competitors is by killing. They have developed two major classes of antagonistic mechanisms to eliminate competitors: diffusible and contact-dependent. Diffusible antibacterial molecules have been extensively described in soil bacteria like *Streptomyces*, which produces antibiotics (e.g. streptomycin, kanamycin, and tetracycline) to kill competitors, gain resources for their own population, and maintain symbiosis with associated plants^2^. *Pseudomonas aeruginosa* is also known to secrete lethal toxins like pyocyanin, exotoxin A, and ExoU that aid in competing against other microbes and human cells during infections^3–5^. On the other hand, contact-dependent antagonisms are less diverse and understudied in the social interaction aspects. The type VI secretion system (T6) discovered in 2006, for example, is a contact-dependent “nano-harpoon” similar to a contractile spear that kills neighboring cells by injecting them with a set of toxic proteins^6^. The T6 is estimated to be found in ~25% of all Gramnegative bacterial species^7^, and targets diverse cell types, including eukaryotes like macrophages and largely Gram-negative bacteria like *Escherichia coli*, in both an environmental and host context^6,8^.

While the regulation, genetics and functional mechanics of the T6 have been well studied^9^, we know relatively little about how targeted cells respond, defend, and survive T6 attack. Similar to antibiotic resistance, one strategy is to neutralize the toxins. Bacteria wielding a T6 that carries anti-microbial toxins do not intoxicate themselves or their sibling cells because a conjugate immunity protein is encoded in the same gene cluster as each toxin^10–12^. However, cells lacking immunity proteins are vulnerable to the toxins. In some cases, bacteria can acquire a library of orphan immunity proteins via horizontal gene transfer and mobile genetic elements, enabling them to survive toxins expressed by unrelated cells^13–16^. *Pseudomonas aeruginosa*, a model organism for T6 research, is able to use cues from the environment to fight back against a T6-wielding aggressor in two ways. In a “tit-for-tat” mechanism, cells that have been intoxicated by T6 can then assemble their own apparatus and launch a counter-attack in the same direction from which the first attack came^17,18^. *P. aeruginosa* is additionally able to induce T6 attack in response to kin cell lysis, via a mechanism called “danger sensing”^19^. Physical processes can also offer protection. Extracellular polysaccharide can protect cells from T6 attack, as does the accumulation of cellular material from lysed cells and physical separation, which are both consequences of T6 antagonism^20–23^. External signaling can play a role in this protection, with recent reports that the presence of glucose enhances survival of *E. coli* cells to T6 attack, mediated through cyclic AMP and its cognate target, the CRP regulator^24^. Other regulators that coordinate stress response systems, such as Rcs and BaeSR may also play an important role, as deletions of these genes reduces survival from attack^25,26^. Transposon sequencing (Tn-seq)^27^ offers one approach to identify genes that affect T6 resistance, uncovering mutations that either increase or decrease survival^28^. However, this technique has a limited range of mutations it can uncover, identifying only single null mutations contributing to a phenotype, but not deleterious mutations in essential genes, functional point mutations, or epistatic relations between multiple genes. Mutagenic screens also do not take pleiotropic sideeffects of mutations into account. For example, mutations that increase T6 survival but come at a steep cost to cellular growth rates would be detected in such a screen, but might not be expected to arise under conditions where reproductive fitness is important.

Experimental evolution^29,30^ circumvents many of these issues, allowing interrogation of the whole genome in a high-throughput, unbiased manner. By including periods of growth between rounds of T6 attack, this approach allows selection to include key pleiotropic fitness effects. Clonal interference among beneficial mutations means that only a small fraction of possible beneficial mutations will arise to high frequency in any given experiment^31^, typically favoring those that are most adaptive. Rather than reporting all possible routes to surviving T6 attack, experimental evolution thus provides insight into genetic mechanisms that provide the largest fitness advantage over hundreds of generations of growth and periodic T6 assault.

In this paper, we explore how *E. coli* evolves resistance to T6 attack by *Vibrio cholerae*. After ~500 generations of growth, punctuated by 30 rounds of attack by the T6, we identified two main mutational pathways, each of which convergently evolved in multiple populations, that enabled dramatically improved survival by *E. coli* during T6 attack. Similar to other types of antibiotic resistance^32^, we find that there was a strong trade-off between increased T6 survival and reduced fitness during growth, which may help explain the continued efficacy of T6 antibiotics in natural populations despite billions of generations of T6 exposure.

## Results

### Experimental evolution of T6 resistance

We report the development of an experimental evolution platform with two model organisms, to identify mechanisms by which bacteria can become resistant to T6 attack (Fig. 1A). We experimentally evolved eight replicate populations of *E. coli* MG1655, exposing them to daily attack by a *V. cholerae* C6706 strain variant that constitutively expresses the building blocks of the apparatus and its four T6 effectors, two that act in the periplasm to degrade the peptidoglycan cell wall (VgrG3 and TseH) and two that disrupt membranes (TseL and VasX) (see Methods and Materials)^12,33–35^. The two species were co-cultured on agar plates in 1:10 ratio (target to killer) to ensure direct contact between cells, which is necessary for T6 attack. 99.99% of our *E. coli* ancestor were killed by *V. cholerae* during the solid-media killing phase of the experiment, imposing strong selection for T6 survival. Between rounds of competition, *E. coli* populations were grown for ~16 generations in LB medium overnight. We also evolved four control populations, competing the same ancestral *E. coli* against a T6-deficient *V. cholerae ΔvasK* strain. We reasoned that mutations arising in these four control populations would account for adaptation in our environment, including growth, dilution, and co-culture with *V. cholerae* on solid media, but not from injury from T6. After 30 rounds of selection, evolved strains were an average of ~27-fold more resistant to *V. cholerae’*s T6 attack, and the control populations had on average 3.9% higher survival, a negligible difference (*F_11,71_* = 15.8, *p* ≤ 0.0001, ANOVA with replicate nested in treatment. Fold survival was log-transformed prior to analysis to homogenize variances, and treatment effect was assessed with pre-planned contrast, *F_1,60_* = 234, *p* ≤ 0.0001; Fig. 1B).

**Figure 1.**
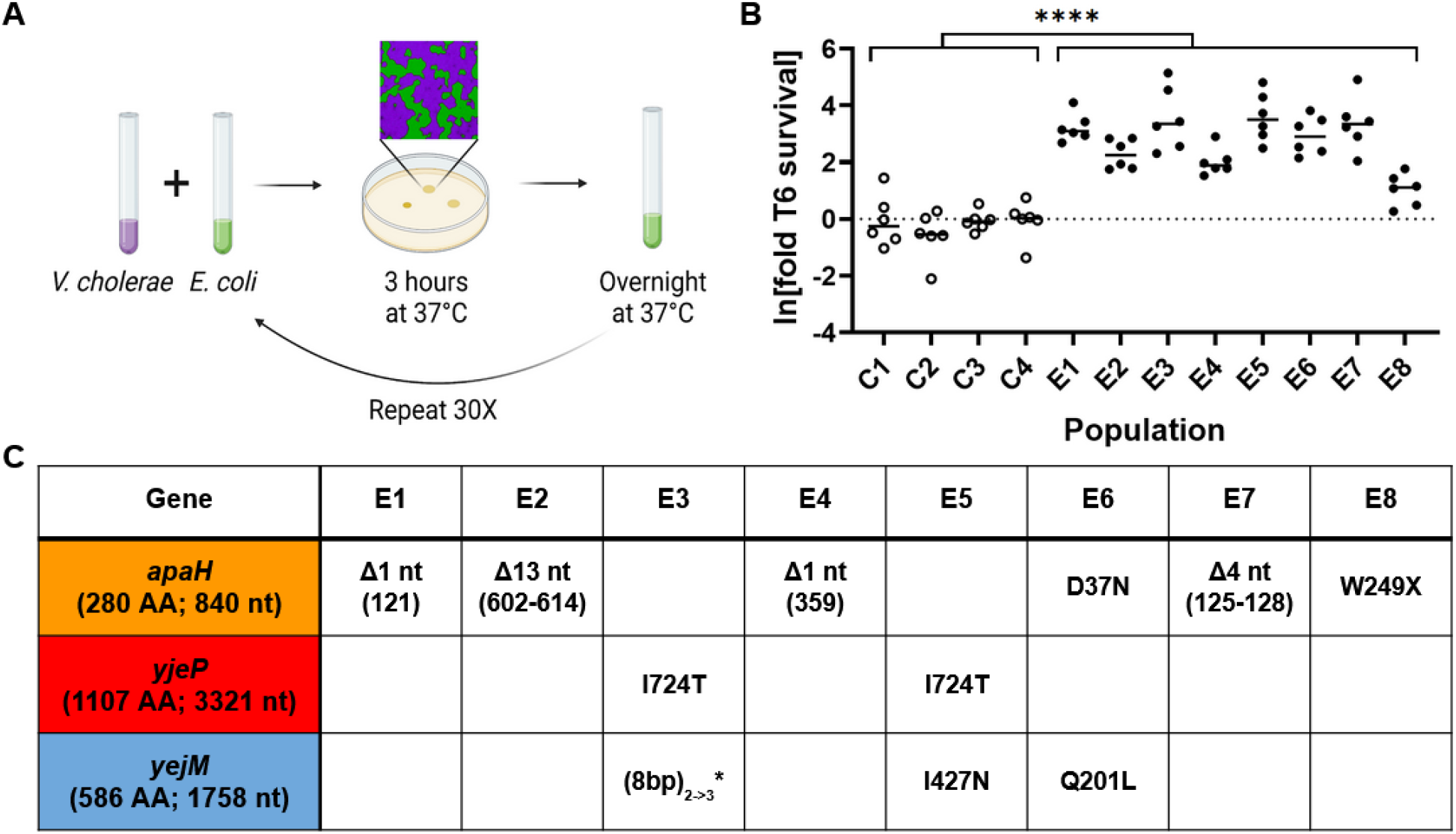
Experimental evolution of resistance to *V. cholerae’*s Type VI Secretion System. (A) Experimental design. We experimentally evolved eight replicate populations of *E. coli*. Each round of selection included ~16 generations of growth in liquid media, followed by co-culture with T6-expressing *V. cholerae* on solid media, where initially the vast majority of *E. coli* were killed. *V. cholerae* were removed via antibiotics, and the surviving *E. coli* resumed growth in liquid media. (B) Over 30 rounds of selection, *E. coli* in the T6 treatment (solid circles) evolved a 27-fold increase in T6 survival, while controls competed against a T6(-) *V. cholerae* (open circles) did not evolve a significant increase in T6 resistance. **** denotes a difference in survival with p□≤□0.0001, determined via ANOVA and a pre-planned contrast. (C) Convergent evolution of genes affording T6 survival. Three genes were mutated in all eight independently evolving populations: *apaH* arose in six, while mutations in *yejM* and *yjeP* arose in the other two populations. For deletions (Δ), numbers in parentheses refer to the nt position of the deletion. (8nt)_2->3_* refers to an 8 nt repeat that expanded from 2 repeats to 3 repeats long, resulting in a frameshift mutation. W249X refers to a premature stop codon at position 249, resulting in a protein product truncated near the C terminus. (AA = amino acids; nt = nucleotides).

### Identifying and characterizing key mutations

We identified mutations arising in our experiment by sequencing a single genotype from each population after 30 rounds of selection. With an average of 2.75 (standard deviation 1.09) mutations per genome in the experimental populations, we chose to focus on mutations that occurred in more than one replicate population, as convergent evolution strongly suggests these mutations are adaptive (Fig. 1C, Fig. S1). Six of the eight isolates had mutations in *apaH*. Four of which are frameshift mutations, suggesting they resulted in loss-of-function (Fig. 1C). This gene is responsible for the “de-capping” of mRNAs in a bacterial cell^36^. Little is known about the global regulatory effect of loss of *apaH*, but it is hypothesized that a null mutation leads to RNA stabilization. Notably, the isolate from population E8 only gained a ~3-fold increase in survival relative to its ancestor; which was significantly lower than five of the seven other replicate experimental populations (Fold survival was log-transformed prior to analysis to homogenize variances, pairwise differences between each replicate population assessed via ANOVA and Tukey’s HSD with overall significance at α = 0.05; Fig. 1B). The mutation in *apaH* found in this isolate creates a premature stop codon near the end of the gene (amino acid 249 out of 280) that likely retains partial function of *apaH*, resulting in a more modest survival advantage.

Two of the eight isolates did not have a mutation in *apaH*. Instead, these two populations each had missense or frameshift mutations in both *yjeP* (also known as *mscM*) and *yejM*, suggesting an interaction between these two genes (Fig. 1C). *yjeP* encodes a mechanosensitive channel that protects cells from osmotic shock^37^. The gene *yejM* (also known as *pbgA* and *lapC*) encodes a metalloprotein that regulates bacterial lipopolysaccharides biosynthesis^38,39^. Deletion of *yejM* is lethal in *E. coli*, while C-terminal truncation mutations result in partial function of the gene^40^. Both mutations we found in *yejM* occur near the C-terminus.

To test the function of mutations found in *apaH, yjeP*, and *yejM* independent of the role of other mutations that arose in experimental lineages (Fig. S1), we re-engineered mutations in these genes in the ancestral strain. A clean deletion of *apaH* increases T6 protection by 3-fold, whereas *E. coli* carrying a single copy of *apaH* expressed from a heterologous constitutive promoter at the Tn7 site is 0.4-fold more susceptible than the ancestor (Fold survival was log-transformed prior to analysis to homogenize variances, comparison of means was accessed with one-sample t-test (μ = 0) and Bonferroni correction with overall significance at α = 0.05, p ≤ 0.0001 and p□≤□0.001; Fig. 2A; see Methods and Materials).

**Figure 2.**
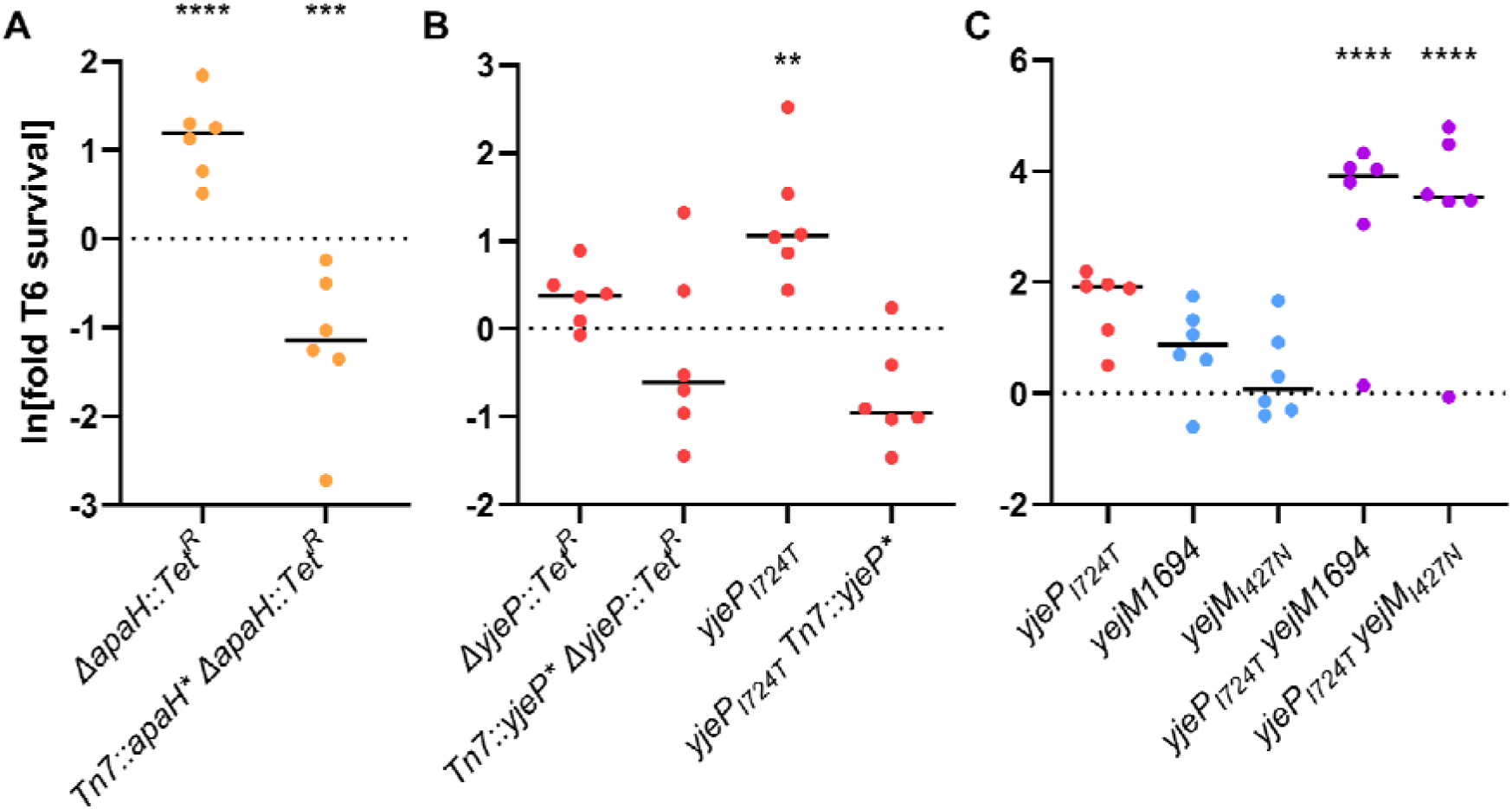
While all mutations of interest increase T6 resistance in various degrees, the *yjeP/yejM* double mutants survive significantly better. (A) *E. coli* with deletion of *apaH* or (B) *yjeP_I724T_* mutation had a slight increase in T6 resistance that was not observed in the other variants. (C) The combination of *yjeP_I724T_* and mutations in the C-terminus of YejM significantly improved the *E. coli* survival by more than 42-fold. Linked markers used to construct the mutants are not indicated in the figure. ****, ***, and ** denote, differences in survival with p□≤□0.0001, pL≤ι_0.001, and p ≤ 0.01 respectively, determined via ANOVA and Dunnett’s Multiple Comparison.

### yjeP_I724T_ is a gain-of-function mutation that confers T6 resistance

*yjeP* is one of four paralogs predicted to encode the MscS mechanosensory channel^37^. An identical missense mutation in *yjeP (yjeP_I724T_*) occurred independently in two lineages (Fig. 1C), suggesting that this amino acid substitution enhances T6 survival and represents a gain-of-function mutation. In the ancestor genetic background, we introduced a *yjeP* disruption, constitutively expressed *yjeP*, and reconstructed the *yjeP_I724T_* mutation. Interestingly, neither the absence of *yjeP* nor its constitutive expression affected T6 survival. However, *E. coli* carrying the *yjeP_I724T_* mutation experienced a ~4-fold survival benefit (Fold survival was log-transformed prior to analysis to homogenize variances, pairwise differences between each replicate population assessed via ANOVA and Dunnett’s test with overall significance at α = 0.05, p≤0.01; Fig. 2B).

Because YjeP is predicted to be a mechanosensitive channel^37^, we determined how the *yjeP_I724T_* mutant responded to pH and osmotic shock, classic stressors for probing mechanosensor function. A *yjeP* null mutant behaved like WT. Interestingly, while the *yjeP_I724T_* mutant *w*as unaffected by changes in osmolarity, it did exhibit 1.1-to 1.4-fold decreases in maximum growth rate in the exponential phase, which was more pronounced with potassium, suggesting that the YjeP may be an ion channel (OD_600_ was log-transformed prior to analysis, and the pH effect was assessed with linear regression comparison of the slopes in the exponential phase with significance at α = 0.05; Fig. 3). To determine whether YjeP is the only MscS mechanosensitive channel protein that can affect T6 resistance, we also tested one of three YjeP homologs, YbdG^37^, because a prior study showed a *ybdG_I176T_* gain-of-function mutation also confers sensitivity to osmotic shock^41^. Unlike *yjeP_I724T_*, the *ybdG_I167T_* did not confer T6 resistance, nor did a *ybdG* null (Fig. 3; Fig. S2). Thus, we conclude that *yjeP_I724T_* is a gain-of-function, or co-dominant, mutation in an ion channel that confers T6 resistance.

**Figure 3.**
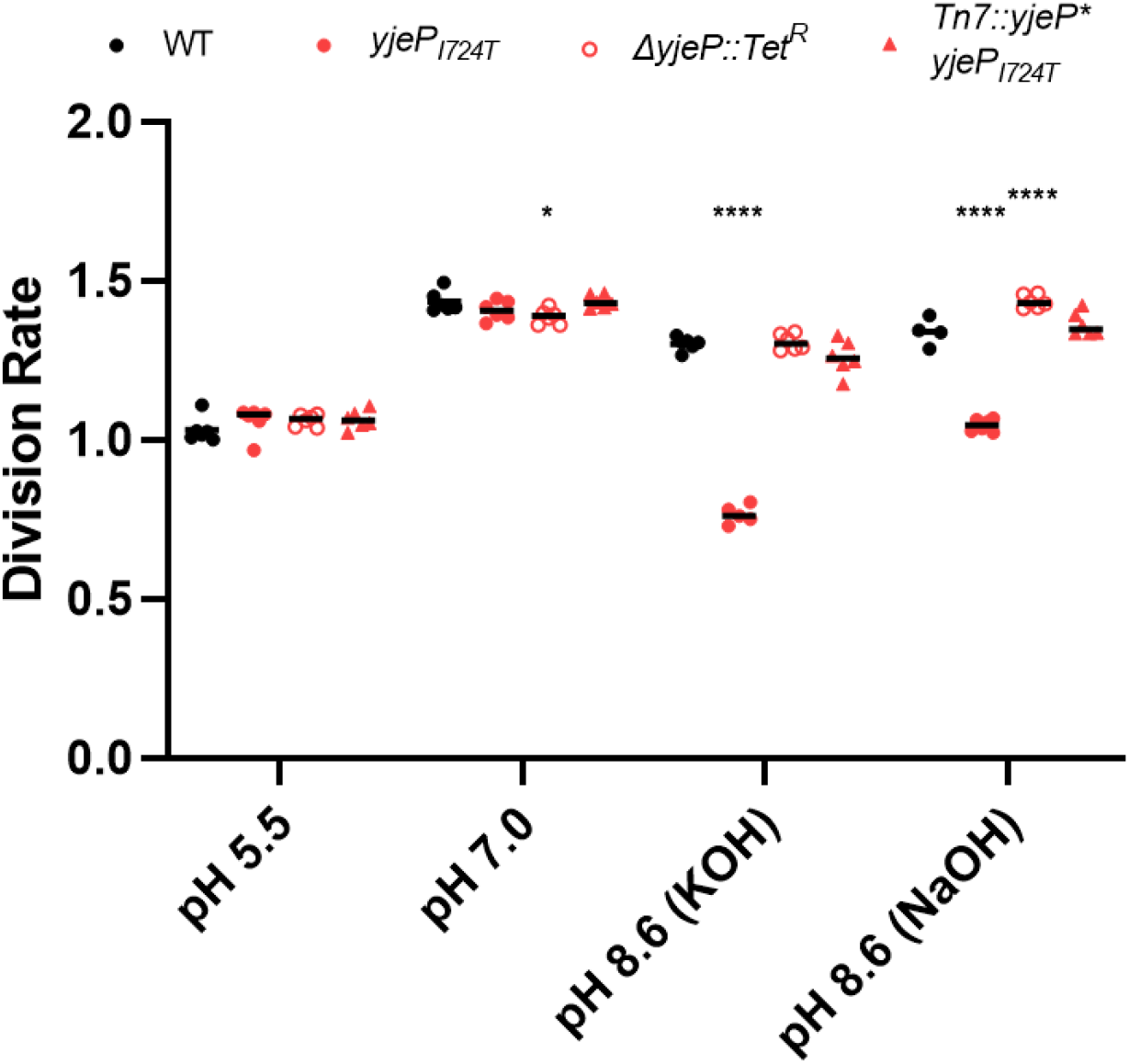
*E. coli yjeP_I724T_* has reduced fitness under basic conditions. *E. coli* and its *yjeP* derivatives grow similarly under acidic and neutral pH. In basic media, however, the *yjeP_I724T_* mutant has a significant decrease in division rate. Linked markers used to construct the mutants do not affect growth in the tested conditions (Fig. S3). ****, **, and * denote, differences in survival with p□≤□0.0001, p□≤□0.01, and p ≤ 0.05 respectively, determined via Two-way ANOVA and Dunnett’s Multiple Comparison.

### E. coli yjeP/yejM double mutants are much more resistant to novel T6 toxins

The fact that *yejM* and *yjeP* accrued mutations in parallel in two independent populations suggests there may be an epistatic relationship between these two mutations. To test this hypothesis, we introduced both *yejM* mutations into the ancestral *E. coli* without and with the *yjeP_I724T_* mutation. While the *yjeP_I724T_* mutation confers a modest benefit (4-fold increased survival; fold survival was log-transformed prior to analysis to homogenize variances, pairwise differences between each replicate population assessed via ANOVA and Dunnett’s test with overall significance at α = 0.05, p≤0.01; Fig. 2B), the presence of either *yejM* mutation by itself has no effect on resistance (Fig. 2C). However, the *yjeP_I724T_* mutation combined with either *yejM* mutation enables a ~40-50-fold increase in survival compared to the ancestor (fold survival was log-transformed prior to analysis to homogenize variances, pairwise differences between each replicate population assessed via ANOVA and Dunnett’s multiple comparison with overall significance at α = 0.05, p≤0.0001; Fig. 2C). In other words, mutation in *yejM* increases resistance only in strains that also have the *yjeP* point mutation.

We next examined whether the mutations that arose in our experiment provide general resistance to T6 attack, or are specific to the toxins employed by the *V. cholerae* C6706 strain, used in this evolution screen, which codes three auxiliary T6 effectors in addition to the large cluster. We therefore competed each mutant *E. coli* strain against an environmental isolate of *V. cholerae* killer, BGT41 (also known as VC22), which encodes a constitutive T6 with effectors predicted to have enzymatic activities distinct from those produced by C6706 and encountered by *E. coli* during experimental evolution^42,43^. This environmental isolate is a superior killer of *E. coli*, relative to C6706^43^, necessitating that we perform our killing assays at a 1:4 killer:target ratio, rather than the 10:1 ratio used with C6706. Evolved strains with *yjeP_I724T_* and *yjeP/yejM* double mutations survived 4,000-fold better than the *E. coli* ancestor, but *apaH* did not measurably increase survival (Fold survival was log-transformed prior to analysis to homogenize variances, pairwise differences between each replicate population assessed via ANOVA and Dunnett’s multiple comparison with overall significance at α = 0.05, p≤0.0001; Fig. 4). In addition, unlike with C6706 killer (Fig. 2C), the *yejM* mutations did not further increase the survival of the *yjeP_I724T_* mutant (Fig. 4). This suggests that I724T in *yjeP* may provide broad spectrum resistance to T6 while protection conferred by mutations in the YejM C-terminus and in *apaH* may depend on the specific effector employed.

**Figure 4.**
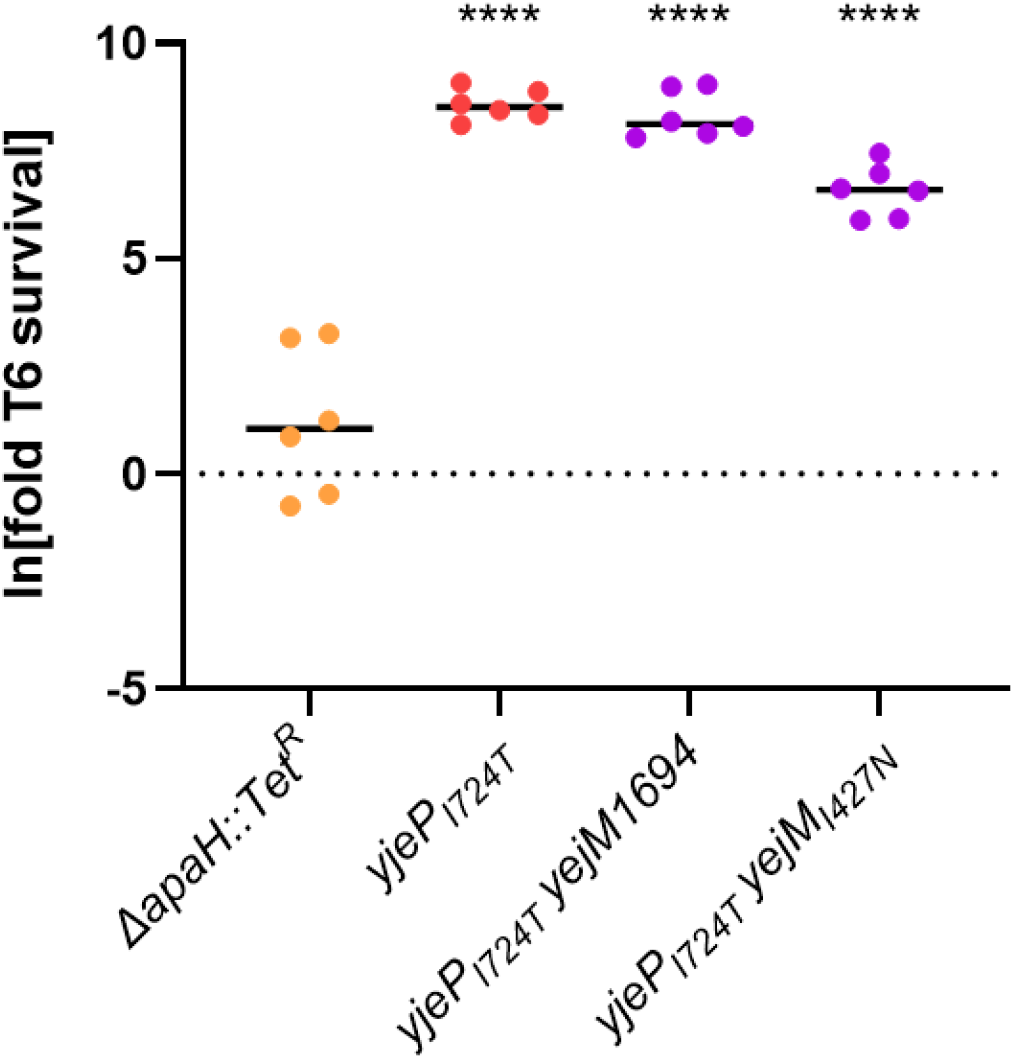
The *yjeP* and the *yjeP/yejM* mutants provide general resistance to T6 attack. When competed against *V. cholerae* with a set of toxins not encountered during experimental evolution, *E. coli* mutants with *yjeP_I724T_* had an increase of >250-fold in T6 resistance whereas deletion of *apaH* did not provide protection against novel T6 effectors. Linked markers used to construct the mutants are not indicated in the figure. **** denotes a difference in survival with p ≤ 0.0001, determined via ANOVA and Dunnett’s Multiple Comparison.

### Experimental evolution reveals trade-offs between T6 resistance and growth rate

So far we have shown that *E. coli* readily evolves resistance to T6 attack, one of the most common mechanisms of antimicrobial warfare. Why, after billions of years of evolution, are bacteria still so poorly defended against T6? Evolutionary theory predicts that trade-offs between antibiotic resistance and other fitness-dependent traits can maintain susceptibility^44^. To test this hypothesis, we examined the effect of each mutation on cellular growth rate by competing them against the ancestral genotype of *E. coli*, under the conditions that mirrored our selection experiment. Mutations in *apaH, yejM*, and *yjeP* decreased fitness during growth (Fig. 5). In fact, there was an overall negative correlation between T6 survival and growth rate for the strains generated in this study (*log_10_(survival) = −2.988 log_10_(growth) −0.2698*, R^2^=0.65, p=3.01*10^-5^; this regression excludes the *crp* and *rlmE* mutants, which never arose during experimental evolution; Fig. 5A).

**Figure 5.**
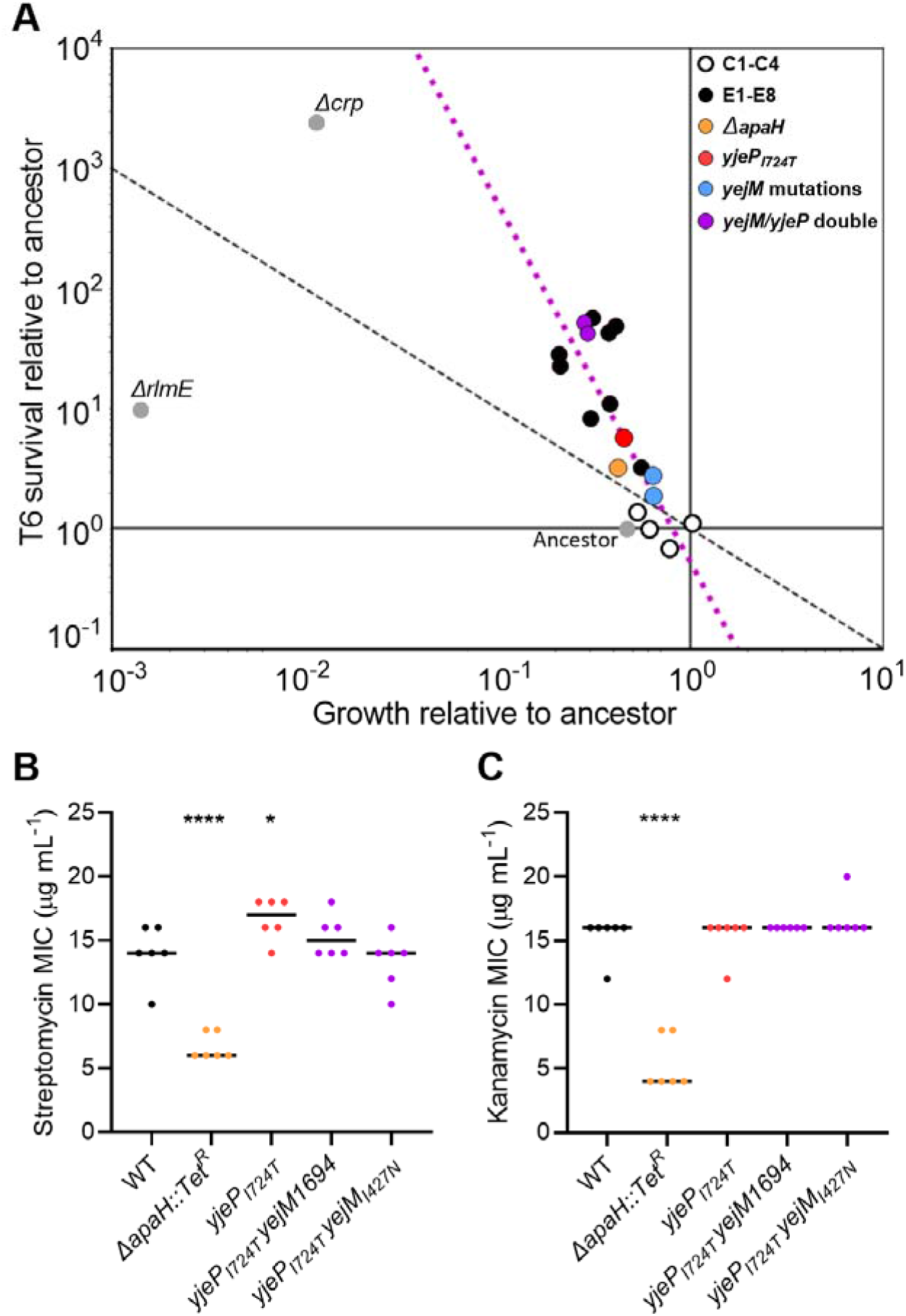
Trade-offs between T6 resistance and fitness during growth. **(A)** Mutations conferring a larger T6 survival advantage also resulted in a greater reduction to reproductive fitness. Plotted are the change in frequency of each mutant across one 16 generation growth assay, and one T6 attack, following the protocols from our evolution experiment. The dashed line represents a fitness isocline, where fitness across one round of selection is equal to that of the ancestor. In other words, the isocline represents where increased fitness during T6 survival is exactly outweighed by decreased fitness in the growth phase. The pink dashed line represents correlation between survival benefit and growth cost; *log_10_(survival) = −2.988 log_10_(growth) - 0.2698*, r^2^=0.65, p=3.01*10^-5^. Green: reconstructed mutations; Red: Evolved isolates; Blue: evolution controls. **(B,C)** Disruption of *apaH* results in decreased MIC for streptomycin (B) and kanamycin (C). The point mutation *yjeP_I724T_* does not affect susceptibility to these antibiotics. Linked markers used to construct the mutants are not indicated in the figure. **** and * denote differences in survival with p ≤ 0.0001 and p□≤ 0.05, determined via ANOVA and Dunnett’s Multiple Comparison.

Our evolution experiment consisted of ~16 generations of exponential growth in LB media, followed by T6 killing. We thus calculated a fitness isocline across the phase space of this trade-off (black dashed line in Fig. 5A), along which a mutant would have equal fitness to the ancestor across one round of growth and killing, with the equation *y* = l/*x* (in log_10_ space). For example, along this line, a 100-fold increase in T6 survival is exactly canceled out by a 100-fold decrease in overnight growth. Mutations that lie above this line should be more fit than our ancestral strain, while mutations below the line should be maladaptive. Perhaps unsurprisingly, given the strong selection on both growth and T6 survival, all mutations we identified are adaptive.

We also measured growth and survival rates for two disruptive mutations that did not arise in our experimentally evolved populations - *crp* and *rlmE*. We have previously shown that deletion of *crp*, a global transcriptional repressor, results in increased survival to the T6 in *E. coli*, but also greatly reduces growth rate^24,45,46^. While this mutation does fall above the fitness isocline, it did not appear in our evolution experiment (Fig. 5). Because all of our mutations of interest result in decreased growth rate, we also sought to test whether decreased growth rate was sufficient to increase T6 resistance. For example, slower growth could prevent microcolonies of the two strains from physical contact on the plate during the course of the coculture competition. We constructed an *E. coli* strain with a *rlmE* deletion, which grows ~0.14% as much as the ancestor during one round of growth (t-test p=8.75*10^-5^). This strain results in much smaller colonies when growing on plates, but has only a 10-fold increase in survival when challenged with T6 attack (Fig. S4). The *rlmE* mutant is below the fitness isocline in Fig. 4A, as the modest increase in survival is not commensurate with the huge growth defect of this mutant if slower growth always led to higher T6 resistance. This shows that slower growth is a sideeffect of mutations that increase T6 resistance, not a cause of increased resistance.

Another trade-off we tested is susceptibility to aminoglycoside antibiotics. The *apaH* disruption strain has a significantly lower minimum inhibitory concentration (MIC) than the ancestor when grown in streptomycin and kanamycin (pairwise differences between each replicate population assessed via ANOVA and Dunnett’s test with overall significance at α = 0.05, p ≤ 0.0001 and p□≤ 0.05; Fig. 5B-C), meaning that they are more susceptible to these antibiotics. This is consistent with previous work on *apaH*^47^. However, strains containing the *yjeP* point mutation did not show increased susceptibility.

## Discussion

In this paper, we use experimental evolution to examine how bacteria adapt to frequent T6 exposure. We subjected populations of *E. coli* to alternating selection for rapid growth followed by attack by *V. cholerae’*s T6 (Fig. 1A). All replicate populations evolving increased T6 resistance (seven of the eight populations) utilized one of two pathways: either a loss-of-function mutation in *apaH;* or a gain-of-function mutation I724T in *yjeP* combined with a partial loss-of-function in *yejM*, with both mutations necessary to provide a large survival advantage (Fig. 1B-C). For a *yjeP_I724T_* mutant, the protection appears to be broad-spectrum, increasing resistance to novel effector proteins by more than 3,000-fold (Fig. 4). Interestingly, the *yjeP/yejM* double mutants are also comparatively resistant to T6 when competing against *V. cholerae* BGT41 (Fig. 4), suggesting *yjeP*_I724T_ provides a broader protection while the additional *yejM* mutations are specific to C6706 T6 effectors (Fig. 2C). While the mechanism underpinning this strain-specific effect is beyond the scope of this study, we hypothesize *yejM_I427N_* still encodes a partially functional YejM periplasmic domain, whereas an insertion of 8 nt (*yejM*1694) results in a frameshift mutation, resulting in a complete disruption of the C-terminus^48^. In *Salmonella enterica and E. coli*, truncation of the C-terminus was shown to disrupt the function of YejM, negatively impacting lipopolysaccharide biosynthesis. This leads to a defective outer membrane that leaks periplasmic proteins into the extracellular space^49^. Periplasmic leakage may reduce the concentration of membrane-localized T6 toxins injected into *E. coli* bearing the *yejM*1694 mutation, reducing their lethality. In contrast, mutations in *apaH* were specific to the T6 effectors they were evolved against, showing no efficacy against a different strain of *V. cholerae* with novel T6 effectors (Figs. 2A, 4).

Of the two primary mutational pathways we focused on in this study, it is interesting that the less beneficial path to T6 resistance, loss of function in *apaH*, evolved more times than the far more beneficial combination of *yjePI724T* and a partial loss-of-function of *yejM* (Fig. 4). This is likely because it is easier to gain beneficial mutations in the *apaH* pathway: any loss of function mutation in the gene gives the phenotype, whereas the *yejM/yejP* pathway requires mutations in two separate genes. The convergent evolution we observed in our experiment (identical *yjeP* SNPs in both populations evolving resistance via this mechanism) further suggests that specific mutations, not simple loss of function mutations, may be required in *yjeP*. Given the difference in T6 resistance between evolved isolates with an *apaH* mutation (Fig. 1B,C) and the constructed *apaH* mutant (Fig. 2A), we hypothesize that other mutations acquired by the evolved populations may also contribute to T6 survival.

Over 500 generations of experimental evolution in 8 replicate populations, we found just two pathways to increased T6 resistance. While prior work has shown that many genes that can affect T6 survival^25,28,50–52^ implying that adaptation might be idiosyncratic among independent populations, our results suggest that adaptive routes to T6 resistance are remarkably constrained. One possibility is that our populations are mutationally limited. This is unlikely, as we can expect ~9.2 x 10^5^ mutations to arise within each growth cycle (based on ~10^10^ cells being produced per cycle, a per base mutation rate of ~(0.2 x 10^-10^)^53^ and a genome size of 4.6 MB), or 2.8 x 10^7^ mutations in each population over the course of the experiment. Instead, the high degree of evolutionary convergence in our experiment suggests that there may simply be relatively few routes to increased T6 survival in which the benefits of the mutation, integrated across the culture cycle to include pleiotropic costs, are great enough to drive the clonal lineage to high frequency.

The evolution of resistance to diffusible antibiotics has been extensively studied^54^. While the details depend on taxon and environment^55,56^, antibiotic resistance often comes with tradeoffs to other fitness components^55–59^. This is especially true for mutations in essential genes that are the target of antibiotics, such as genes encoding ribosomal proteins^60^. However, compensatory evolution often reduces initially severe costs of resistance, either via the fixation of epistatic mutations elsewhere in the genome, or by replacing initially costly resistance mutations with lower-cost alternatives^60–63^. In contrast to diffusible antibiotics, the eco-evolutionary dynamics of contact-mediated killing remains largely unexplored, and it is unclear if or when similar compensatory adaptation would occur if we continued our experiment. The fact that we observe a strong trade-off between T6 survival and growth rate is not entirely unexpected. The T6 secretion system is an ancient, widespread, and highly effective microbial weapon. Trade-off free adaptations that increase survival to T6 attack would be expected to rapidly fix in many bacterial populations. As a result, pleiotropic costs to T6 resistance could play an important role in maintaining T6 efficacy over evolutionary time.

Single mutations that confer resistance to an individual antibiotic are common, as a modification of one target site may be sufficient to escape drug toxicity^64^. Because the probability a susceptible cell will simultaneously gain mutations allowing it to survive multiple antibiotics is far lower than the probability of gaining resistance to any single antibiotic^65^, physiological mechanisms that afford broad-spectrum toxin resistance (e.g., efflux pumps) can often incur fitness trade-offs^66^. Current efforts to combat antibiotic resistance appropriately focus on identifying drug targets that incur large fitness costs; with modern drug combination, drug cycling, and adaptive therapies seeking to exploit these fitness trade-offs to slow the rate of resistance evolution^67–70^. We thus might expect that, as in our experiment here, T6 resistance often evolves via mechanisms that modify cellular physiology or behavior (e.g., increased capsule thickness) that improves survival, albeit with pleiotropic costs^21^. In contrast to diffusible antibiotics, it may be more difficult for bacteria to evolve resistance to T6-delivered antibiotics. T6 attacks synchronously deliver multiple effectors that target different components of the intoxicated cell, and delivery is direct, which minimizes dilution and dispersal of the toxins in a heterogenous extracellular environment.

It has only recently become apparent how important social interactions are to microbial ecology and evolution^71–73^. Antagonistic interactions appear to be more common than cooperation or commensalism^1^, at least for species that are capable of being cultured. The Type VI secretion system - a ballistic harpoon containing multiple types of toxins capable of quickly killing susceptible cells, represents the cutting-edge of microbial weaponry. In this paper, we show that *E. coli* can indeed evolve substantial genetic resistance to T6 assault, but doing so entails trade-offs with reproductive fitness. We also found that one convergently-evolving solution appeared to provide effector-specific protection, while the other appeared to be more general. So far, relatively little effort has gone into understanding the mechanisms (both genetic and behavioral) through which microbes can evolve to resist dying from T6- a crucial gap in our knowledge that limits our ability to understand the ecology and evolution of this widespread microbial weapon. Further work will be required to determine if trade-offs between T6 survival and reproduction are found in other taxa, and whether or not such trade-offs can be mitigated over longer evolutionary timescales via compensatory mutation^74^.

## Materials and Methods

### Bacterial strains and media

Bacterial strains were grown aerobically at 37 °C overnight in lysogeny broth (LB) (1% w/v tryptone (Teknova, CA, USA), 0.5% w/v yeast extract (Hardy Diagnostics, CA, USA), 1% w/v NaCl (VWR Life Sciences, PA, USA) or liquid basal medium (100 mM Tricine (Thermo Scientific, MA, USA), 10 mM K_2_HPO_4_ (Fisher Scientific, NH, USA), 0.5% w/v tryptone, 0.25% w/v yeast extract, 0.5% w/v glucose (VWR, PA, USA), and pH 5.5 with HCl (Fisher Scientific, NH, USA) or pH 8.6 with KOH (Fisher Scientific, NH, USA) or NaOH (Fisher Scientific, NH, USA)) with constant shaking or on LB agar (1.5% w/v agar; Genesee Scientific and Hardy Diagnostics, CA, USA). Ampicillin (GoldBio, MO, USA and VWR Life Sciences, PA, USA), spectinomycin (Sigma-Aldrich, MO, USA and Enzo Life Sciences, NY, USA), streptomycin (VWR Life Sciences, PA, USA), kanamycin (GoldBio, MO, USA and VWR Life Sciences, PA, USA), chloramphenicol (Sigma-Aldrich, MO, USA and EMD Millipore, MA, USA), tetracycline (Sigma-Aldrich, MO, USA and Fisher BioReagents, PA, USA), and arabinose (GoldBio, MO, USA) were supplemented where appropriate. Specific concentrations will be described below.

### Mutant construction

Mutations were introduced into *E. coli* K-12 strain MG1655 *ΔaraBAD::cat* by P1vir transduction^75^. Point mutations in *ybdG, yejM*, and *yjeP* were transduced into the recipient strain using the genetically linked markers *purE*79::Tn10, zei-722::Tn10, and *ΔyjeJ::ampR*, respectively. Transductants were selected for using 10 μg mL^-1^ tetracycline or 25 μg mL^-1^ ampicillin and screened for the presence of the point mutations by DNA sequencing (Azenta Life Sciences, MA, USA). All null mutations were confirmed by PCR.

*yjeJ* and *rlmE* were deleted and replaced with the Amp^R^ or Tet^R^ cassette, respectively, by λ Red recombination as previously described^76^. To generate *ΔyjeJ::ampR*, the Amp^R^ cassette from pUC19 was amplified by PCR using the primers KOyjeJBla.Fwd and KOyjeJBla.Rev, which contain homology to the 5’ and 3’ ends of *yjeJ*, respectively. To generate *ΔrlmE::tetA*, the *tetA* gene and promoter were amplified from Tn10 using the primers rrmJTET.Fwd and rrmJTET.Rev. *ΔyjeJ::ampR* or *ΔrlmE::tetA* DNA were transformed into DY378, a strain of *E. coli* K-12 that expresses the λ Red recombination system from a temperature sensitive promoter. Prior to transformation, the λ Red system was induced by incubating midlog phase DY378 cells at 42°C for 15 minutes in a shaking water bath. Recombinants were selected for on LB containing 25 μg mL^-1^ ampicillin (for *ΔyjeJ::ampR*) or 10 μg mL^-1^ tetracycline (for Δ*rlmE*::*tetA*).

To generate the *ΔapaH::tetA, ΔybdG::tetA*, and *ΔyjeP::tetA* null alleles, *ΔapaH::ampR, ΔybdG::kanR*, and *ΔyjeP::kanR* from the Keio library^77^ were moved into DY378 by P1vir transduction^75^. The Kan^R^ cassette in each Keio allele was replaced with *tetA* from Tn10 by λ Red recombination^76^. The *tetA* DNA was amplified by PCR using the primers pKD13TetA.Fwd and pKD13TetA.Rev, which contain homology to the 5’ and 3’ ends of the Kan^R^ cassette, respectively. Recombinants were selected for on LB containing 10 μg mL^-1^ tetracycline and screened for sensitivity to 25 μg mL^-1^ kanamycin.

*ybdG*_I167T_ was constructed using CRISPR-Cas9 gene editing as previously described^78^. The *ybdG* guide RNA plasmid pCRISPR-ybdG493 was constructed by ligating ybdG493.CRISPR duplexed DNA (Integrated DNA Technologies, IA, USA) into BsaI-digested pCRISPR. 100 ng of pCRISPR-ybdG493 and 10 uM of the editing oligonucleotide ybdGI167T.MAGE (Integrated DNA Technologies, IA, USA) were transformed into MG3686, a derivative of DY378 that constitutively expresses Cas9 from a plasmid. Transformants were selected for on LB containing 25 μg mL^-1^ chloramphenicol and 50 μg mL^-1^ kanamycin. Recombinants containing the *ybdG*_I167T_ mutation were identified by DNA sequencing (Azenta Life Sciences, MA, USA). Two phosphorothioate bonds were added at the 5’ and 3’ ends of the ybdGI167T.MAGE oligonucleotide to increase stability.

Genes were inserted at the Tn7 attachment site following a similar protocol described previously^79,80^. Wildtype *apaH* or *yjeP* expressed from the constitutive promoter J23119 (http://parts.igem.org/Part:BBa_J23119) were cloned into XhoI and HindIII (New England Biolabs, MA, USA) digested pZS21, resulting in the plasmids pZS21-*apaH* and pZS21-*yjeP*. The J23119 promoter, gene, and *rrnB1* terminator from pZS21-*apaH* or pZS21-*yjeP* were amplified by PCR using the primers pGRG25GA.Fwd and pGRG25GA.Rev. The Ω streptomycin/spectinomycin resistance cassette from pHP45Ω was amplified using the primers pGRG25SpcGA.Fwd and pGRG25SpcGA.Rev. *apaH* or *yjeP* DNA along with DNA corresponding to the Ω streptomycin/spectinomycin resistance cassette were inserted into PacI and AvrII digested pGRG25-ModularBamA-Kan by Gibson Assembly (New England Biolabs, MA, USA). The resulting plasmids were transformed into MG1655 and transformants were selected for on LB containing 25 μg mL^-1^ spectinomycin and 0.2% (w/v) arabinose. Transformants were screened for integration of *apaH* or *yjeP* and the Ω streptomycin/spectinomycin resistance cassette at the Tn7 site by PCR.

*V. cholerae* was genetically engineered with established allelic exchange methods using vector pKAS32^81^. Expression of chromosomal *qstR* from a heterologous Ptac promoter results in constitutive T6 activity because C6706 lacks a functional *lacI* gene^82^. An in-frame deletion of *vasK* tprevents T6 assembly, as described previously^13^. All Insertions, deletions, and mutations were confirmed by PCR and DNA sequencing (Eton Bioscience Inc, NC, USA).

### Experimental evolution

Twelve replicate populations of *E. coli* with chloramphenicol (10 μg mL^-1^) were initiated from an overnight culture of MG1655 with chromosomal Cm^R^ cassette and a plasmid encoding Kan^R^ cassette. Each round, cultures were washed twice with LB to remove antibiotics, then mixed with an overnight culture of either *V. cholerae* C6706 *qstR\** (for the 8 experimental populations) or C6706 *qstR*ΔvasK* (for the 4 control populations) in a 10:1 killer to target ratio. 50 μL of each mixture was spotted onto an LB agar plate, dried, and incubated at 37°C for 3 hours. Competition mixtures were then resuspended in 5 mL of ddH_2_O containing kanamycin (50 μg mL^-1^) and chloramphenicol (10 μg mL^-1^), and put at 4°C for 30 minutes, conditions which allow for survival of *E. coli* but not *V. cholerae*. Surviving cells were then diluted 10-fold into LB containing kanamycin (50 μg mL^-1^) and chloramphenicol (10 μg mL^-1^) for overnight growth at 37°C. This procedure was repeated daily for 30 rounds. A sample of each whole population was frozen at −80°C every five rounds. At the end of 30 rounds, a clonal isolate from each population was taken for subsequent phenotypic and genomic testing.

### Stress assay

The optical density (OD_600_) of overnight cultures of *E. coli* strains growing in the basal medium (pH 7) was measured by a ThermoFisher Scientific Genesys 30 spectrophotometer (MA, USA) and normalized to 1. Then cells were diluted 1:50 into the basal medium (pH 5.5, pH 7, and pH 8.6 with KOH or NaOH) in a 96-microtiter plate, which was incubated aerobically at 37 °C with shaking in a BioTek Synergy H1 microplate reader (VT, USA). The OD_600_ of each well was read every 30 mins for 16 h. A curve of best fit was assigned to each well using the 4P Growth model in JMP (JMP®, Version 16.1.0. SAS Institute Inc., Cary, NC, 1989–2021), and the value of the “Division” parameter was compared across treatments and replicates.

### Antibiotic minimum inhibitory concentration (MIC) determination

Antibiotics were added to wells of a 96-microtiter plate, starting at 1280 μg mL^-1^ for streptomycin and 640 μg mL^-1^ for kanamycin, and serially diluted 2-fold across the plate. Overnight cultures of bacteria were diluted and added to the wells for a final OD_600_ of 0.001. Once a target range was determined to contain the MIC for each antibiotic, a linear range of antibiotic concentrations were prepared and tested in 96-microtiter plate (4 through 36 μg mL^-1^ for kanamycin and 2 through 18 μg mL^-1^ for streptomycin), and bacteria were added at a an OD_600_ of 0.001. Plates were incubated stationary at 37°C for 24 hours. A well was determined to have growth if the OD_600_ was above 0.2, as measured by a BioTek Synergy H1 microplate reader (VT, USA), and the MIC was determined to be the lowest concentration at which no growth occurred.

### T6-mediated competition assay

The OD_600_ of overnight cultures of *V. cholerae* killer and Cm^R^ *E. coli* target were normalized to 1. Killer and target are then mixed in either 10:1 or 1:4 ratio, inoculated onto a pre-dried LB agar, and allowed to dry. After 3 hours of static incubation at 37°C, cells were resuspended in 5 ml of LB, following with serial dilutions. Finally, the resuspension was inoculated on a LB agar containing chloramphenicol (10 μg mL^-1^) to select for the surviving *E. coli*, which was incubated overnight at 37 °C and the *E. coli* colonies were counted. Data is presented as the fold increase of the survival rate for a given genotype as compared to the ancestor (measured in the same experiment), as given by:

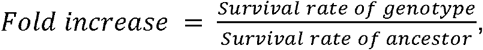

where the survival rate for each strain is calculated by dividing recovered *E. coli* colonies from competition with the T6+ *V. cholerae* strain by the number of colonies recovered from competition with the T6-strain.

### Genomic DNA preparation, whole genome sequencing, and genomic analysis

*E. coli* genomic DNA from each population was isolated using Promega Wizard Genomic DNA Purification Kit (Madison, WI). The DNA quality was analzyed using gel electrophoresis to confirm no degradation and NanoDrop to confirm the purity of the samples (260/280 = 1.8-2.0). Whole genome sequencing was conducted using Illumina sequencing on a NextSeq 2000 platform at Microbial Genome Sequencing Center (PA, USA). Once we received the DNA sequencing results, quality check, filter, base correction, adapter trimming, and merging were conducted using fastp v0.20.0^83^. Reads were then mapped and compared to the *E. coli* MG1655 reference genome (accession U00096) from NCBI Genome database using Breseq v0.35.1 with bowtie2-stage2^84–86^. Other parameters remain default.

## Acknowledgments and funding sources

Figure 1A was made using Biorender.com. K.M. is supported by NIH T32 grant T32GM142616. SLN is supported by NSF grant BMAT 2003721. T.J.S. is supported by NIGMS MIRA grant 5R35GM118024. The content is solely the responsibility of the authors and does not necessarily represent the official views of the National Institutes of Health.

We declare no conflict of interest.

## Data Availability

All data from this study are included in the article and/or supporting information.

