## Supplemental material for "Trade-offs constrain adaptive pathways to T6 survival"

### Supplemental Materials

**Predicted mutations**

| position | mutation | C1-ref | C2-ref | C3-ref | C4_ref | E1-ref | E2-ref | E3-ref | E4-ref | E5-ref | E6-ref | E7-ref | E8-ref | anc-ref | annotation | gene | description |
|---|---|---|---|---|---|---|---|---|---|---|---|---|---|---|---|---|---|
| 50,476 | C→T | | | | | | | | | | | | 100% | | W249* (TGG→TGA) | *apaH* ← | diadenosine tetraphosphatase |
| 50,609 | Δ13 bp | | | | | | 100% | | | | | | | | coding (602-614/843 nt) | *apaH* ← | diadenosine tetraphosphatase |
| 50,864 | Δ1 bp | | | | | | | | 100% | | | | | | coding (359/843 nt) | *apaH* ← | diadenosine tetraphosphatase |
| 51,095 | Δ4 bp | | | | | | | | | | | 100% | | | coding (125-128/843 nt) | *apaH* ← | diadenosine tetraphosphatase |
| 51,102 | Δ1 bp | | | | | 100% | | | | | | | | | coding (121/843 nt) | *apaH* ← | diadenosine tetraphosphatase |
| 51,114 | C→T | | | | | | | | | | 100% | | | | D37N (GAT→AAT) | *apaH* ← | diadenosine tetraphosphatase |
| 134,935 | IS*1* (–) +9 bp | | | | | 100% | | | | | | | | | coding (640-648/795 nt) | *speD* ← | S-adenosylmethionine decarboxylase proenzyme |
| 257,908 | Δ776 bp | 100% | 100% | 100% | 100% | 100% | 100% | 100% | 100% | 100% | 100% | 100% | 100% | 100% | | *insB9*–*[crl]* | *insB9*, *insA9*, *[crl]* |
| 366,351 | 2 bp→TT | 100% | 100% | | | | 100% | | | | | | 100% | | intergenic (-46/+76) | *lacZ* ← / ← *lacI* | beta-galactosidase/DNA-binding transcriptional repressor LacI |
| 367,573 | G→A | 100% | 100% | 100% | 100% | 100% | 100% | 100% | 100% | 100% | 100% | 100% | 100% | 100% | intergenic (-63/+14) | *lacI* ← / ← *mhpR* | DNA-binding transcriptional repressor LacI/DNA-binding transcriptional activator MhpR |
| position | mutation | C1-ref | C2-ref | C3-ref | C4_ref | E1-ref | E2-ref | E3-ref | E4-ref | E5-ref | E6-ref | E7-ref | E8-ref | anc-ref | annotation | gene | description |
| 967,549 | G→T | | | | | 100% | | | | | | | | | R310L (CGC→CTC) | *msbA* → | ATP-binding lipopolysaccharide transport protein |
| 1,074,056 | G→T | | | | 100% | | | | | | | | | | intergenic (-45/-186) | *rutA* ← / → *rutR* | pyrimidine oxygenase/DNA-binding transcriptional dual regulator RutR |
| 1,212,337 | A→G | | 100% | | | | | | | | | | | | intergenic (-33/+203) | *ymgK* ← / ← *ymgL* | protein YmgK/protein YmgL |
| 1,261,524 | A→C | | | | | | | | | | 100% | | | | S118A (TCC→GCC) | *prs* ← | ribose-phosphate diphosphokinase |
| 1,267,961 | T→A | | | | | | | | | | 100% | | | | D214E (GAT→GAA) | *ychA* → | transglutaminase-like/TPR repeat-containing protein |
| 1,299,499 | Δ1,199 bp | ? | ? | ? | ? | ? | ? | 100% | ? | ? | ? | 100% | ? | ? | | *insH21* | insH21 |
| 1,637,786 | IS*1* (+) +9 bp | | 100% | | | | | | | | | | | | intergenic (-4/+160) | *gnsB* ← / ← *ynfN* | Qin prophage; protein GnsB/Qin prophage; protein YnfN |
| 1,705,932 | G→A | | | | | | | | 100% | | | | | | A56T (GCC→ACC) | *rsxA* → | SoxR [2Fe-2S] reducing system protein RsxA |
| 1,978,503 | Δ776 bp | 100% | 100% | 100% | 100% | 100% | 100% | 100% | 100% | 100% | 100% | 100% | 100% | 100% | | *insB-5*–*insA-5* | *insB-5*, *insA-5* |
| 2,173,363 | Δ2 bp | 100% | 100% | 100% | 100% | 100% | 100% | 100% | 100% | 100% | 100% | 100% | 100% | 100% | pseudogene (915-916/1358 nt) | *gatC* ← | galactitol-specific PTS enzyme IIC component |
| position | mutation | C1-ref | C2-ref | C3-ref | C4_ref | E1-ref | E2-ref | E3-ref | E4-ref | E5-ref | E6-ref | E7-ref | E8-ref | anc-ref | annotation | gene | description |
| 2,285,655 | T→A | | | | | | | | | 100% | | | | | I427N (ATC→AAC) | *yejM* → | putative cardiolipin transport protein |
| 2,286,069 | $(GTGAAAGA)_{2\rightarrow3}$ | | | | | | | 100% | | | | | | | coding (1694/1761 nt) | *yejM* → | putative cardiolipin transport protein |
| 2,521,007 | Δ120 bp | | 100% | 100% | 100% | 100% | 100% | 100% | 100% | | | 100% | 100% | | | *[valX]* | *[valX]* |
| 2,561,053 | C→T | | | | | | | | | | 100% | | | | intergenic (+155/-315) | *yffL* → / → *yffM* | CPZ-55 prophage; uncharacterized protein YffL/CPZ-55 prophage; uncharacterized protein YffM |
| 2,912,166 | A→G | | 100% | | | | | | | | | | | | F496L (TTC→CTC) | *relA* ← | GDP/GTP pyrophosphokinase |
| 3,560,455 | +G | 100% | 100% | 100% | 100% | 100% | 100% | 100% | 100% | 100% | 100% | 100% | 100% | 100% | pseudogene (151/758 nt) | *glpR* ← | DNA-binding transcriptional repressor GlpR |
| 3,913,505 | A→T | | | | | | | | | | | | 100% | | L55Q (CTG→CAG) | *glmS* ← | L-glutamine--D-fructose-6-phosphate aminotransferase |
| 4,051,187 | IS2 (+) +5 bp | | | | | | | | | | | 100% | | | noncoding (152-156/245 nt) | *csrC* → | small regulatory RNA CsrC |
| 4,213,066 | C→A | | | | | | | | | | | 100% | | | noncoding (27/120 nt) | *rrfE* → | 5S ribosomal RNA |
| 4,296,381 | +GC | 100% | 100% | 100% | 100% | 100% | 100% | 100% | 100% | 100% | 100% | 100% | 100% | 100% | intergenic (+587/+55) | *gltP* → / ← *yjcO* | glutamate/aspartate : H(+) symporter GltP/Sel1 repeat-containing protein YjcO |
| position | mutation | C1-ref | C2-ref | C3-ref | C4_ref | E1-ref | E2-ref | E3-ref | E4-ref | E5-ref | E6-ref | E7-ref | E8-ref | anc-ref | annotation | gene | description |
| 4,353,486 | IS2 (+) +5 bp | | | | | | 100% | | | | | | | | coding (1228-1232/1518 nt) | *lysU* ← | lysine--tRNA ligase/Ap4A synthetase/Ap3A synthetase |
| 4,353,655 | IS5 (+) +4 bp | | | | | | | | | | | 100% | | | coding (1060-1063/1518 nt) | *lysU* ← | lysine--tRNA ligase/Ap4A synthetase/Ap3A synthetase |
| 4,358,274 | IS5 (+) +4 bp | 100% | | | | | | | | | | | | | coding (341-344/2148 nt) | *cadA* ← | lysine decarboxylase 1 |
| 4,387,200 | A→G | | | | | | | 100% | | 100% | | | | | I724T (ATC→ACC) | *mscM* ← | miniconductance mechanosensitive channel MscM |
| 4,388,769 | T→A | | | | | | | | | | 100% | | | | Q201L (CAG→CTG) | *mscM* ← | miniconductance mechanosensitive channel MscM |

**Figure S1. Breseq result summary.** Genome sequences were compared against *E. coli* MG1655 reference genome (accession U00096) using Breseq. “100%” indicates there are significant differences at the specific position between the compared genomes. “?” indicates where the coverage at the specific position was too low to call that a mutation. Description shows the annotated function or feature of the genes based on the reference genome.

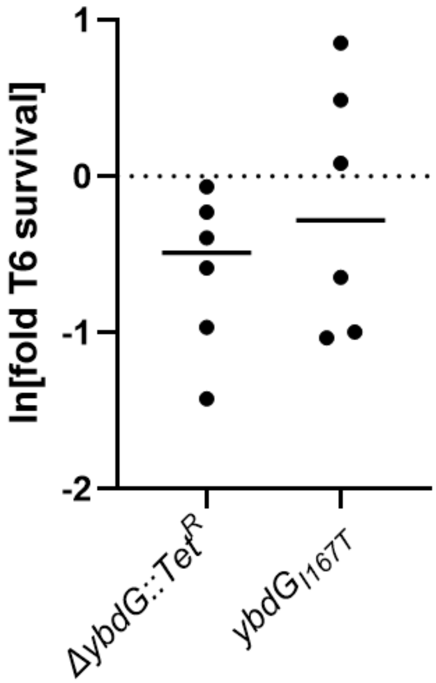


**Figure S2. Gain of Function Mutation in YjeP homolog YbdG does not affect T6 survival in *E. coli*.** Data shows no significant difference in T6 survival when comparing the *ybdG* mutants to the WT *E. coli*. Linked markers used to construct the mutants are not indicated in the figure.

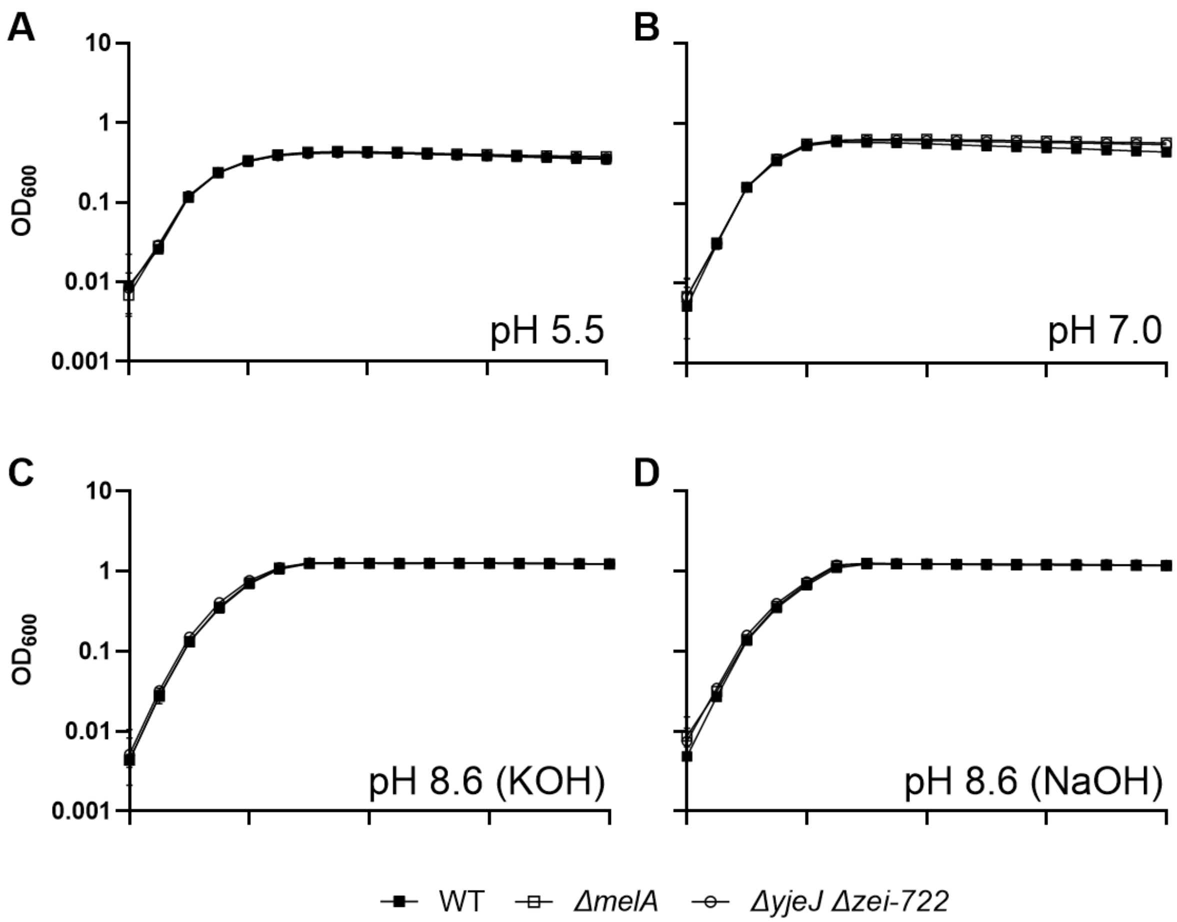


**Figure S3. Linked markers used to introduce mutations in *E. coli* do not affect the growth in tested conditions.** Linkers introduced into *E. coli* did not have significant growth differences in the tested pH concentrations.

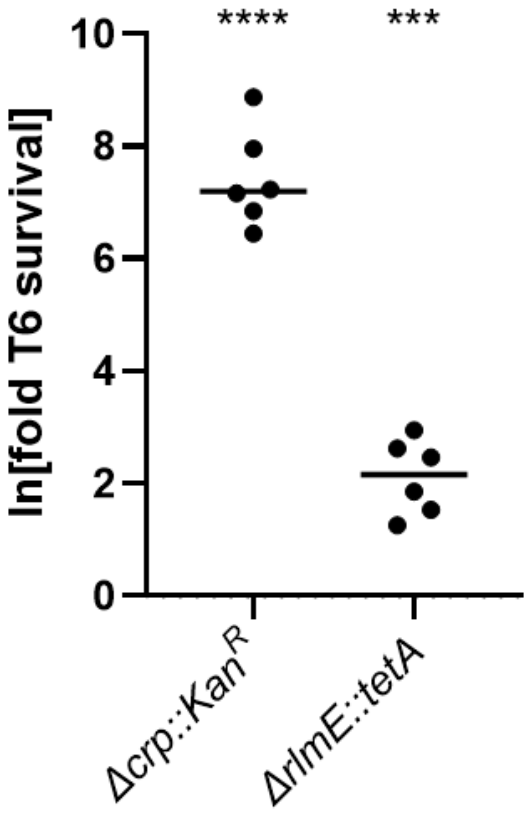


**Figure S4. T6 survival of slow growing *E. coli*.** A *crp* null mutant survives T6 attack over 2,000-fold better than the ancestor, while an *rmlE* mutant only survives ~10-fold better.
